# Adult fibroblasts use aggresomes only in distinct cell-states

**DOI:** 10.1101/2021.07.23.453501

**Authors:** Christopher S. Morrow, Zachary P. Arndt, Bo Peng, Eden Y. Zewdie, Bérénice A. Benayoun, Darcie L. Moore

## Abstract

The aggresome is a protein turnover system in which proteins are trafficked along microtubules to the centrosome for degradation. Despite extensive focus on aggresomes in immortalized cell lines, it remains unclear if the aggresome is conserved in all primary cells and all cell-states. Here we examined the aggresome in primary adult mouse dermal fibroblasts shifted into four distinct cell-states. We found that in response to proteasome inhibition, quiescent and immortalized fibroblasts formed aggresomes whereas proliferating and senescent fibroblasts did not. Using this model, we generated a resource to better understand the aggresome, providing a characterization of the proteostasis networks in which the aggresome is used and transcriptomic features associated with the presence or absence of aggresome formation. Using this resource, we validate a previously reported role for p38 MAPK signaling in aggresome formation and identify TAK1 as a novel driver of aggresome formation upstream of p38 MAPKs. Together, our data demonstrate that the aggresome is a non-universal protein degradation system which can be used cell-state specifically and provide a resource for better understanding aggresome formation and function.

## Introduction

Proper cellular function relies upon effective maintenance of the proteome by the cell’s protein quality control systems, such as the ubiquitin proteasome system, autophagy, and chaperone proteins (Vilchez, Saez et al. 2014, Sontag, Samant et al. 2017, Johnston and Samant 2020). In response to environmental challenges which impair proteostasis, cells react by regulating each of these pathways to regain homeostasis. Additionally, many cell types form an aggresome, a structure comprised of proteins destined for degradation being trafficked along microtubules by dynein and other adapter proteins to the centrosome, within a cage comprised of the intermediate filament vimentin (Fig. 1A) (Johnston, Ward et al. 1998, Kopito 2000, Johnston, Illing et al. 2002, Iwata, Riley et al. 2005, Olzmann, Li et al. 2008). It is thought that aggresome formation is cytoprotective by increasing the efficiency of protein degradation by bringing together proteins destined for degradation with protein degradation machineries and organizing damaged or misfolded proteins so they do not interfere with other cellular processes (Tanaka, Kim et al. 2004, Olzmann, Li et al. 2008, Morrow, Porter et al. 2020, Sunchu, Riordan et al. 2020).

**Figure 1.**
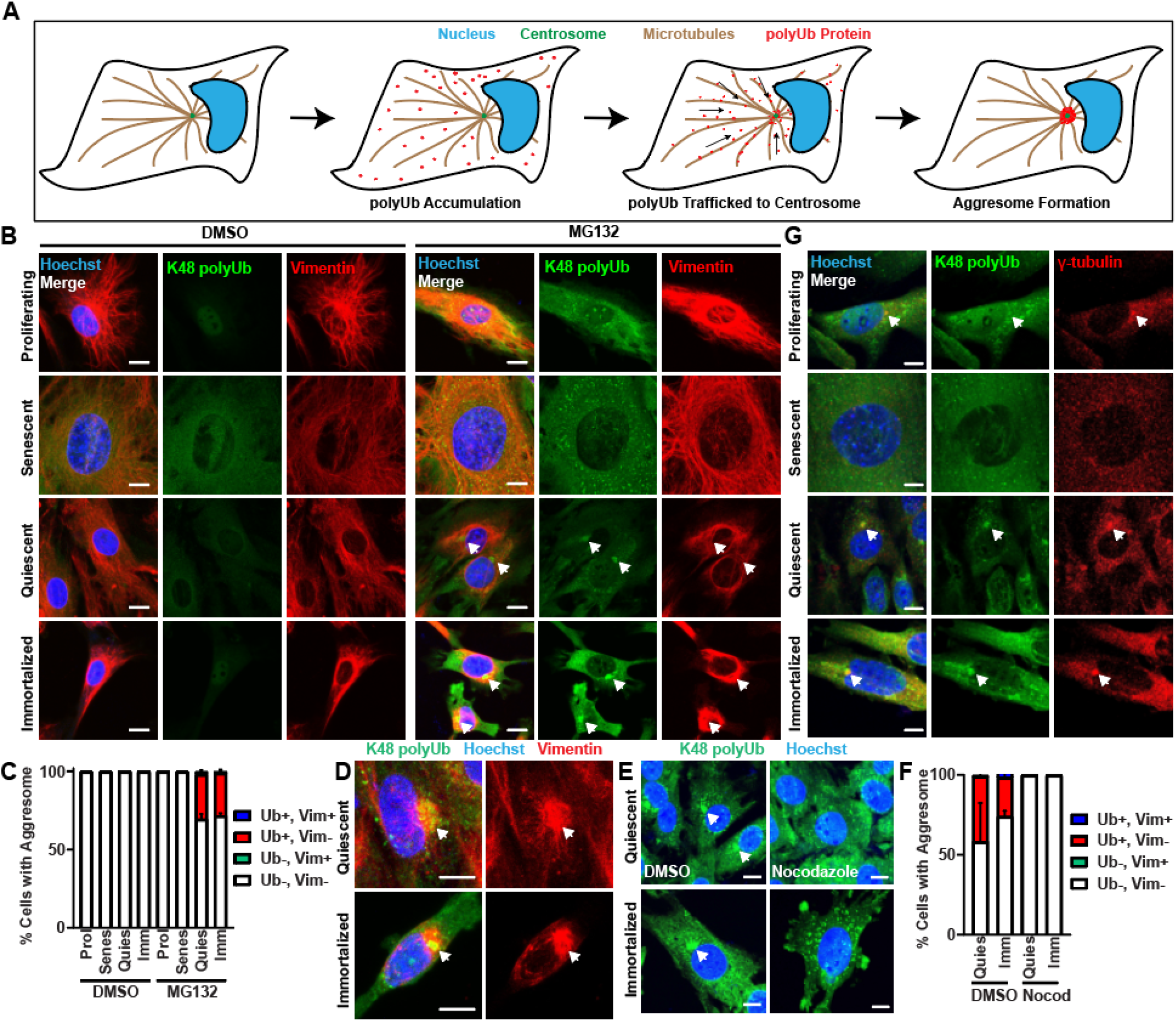
Aggresomes are formed by fibroblasts in response to proteasome inhibition in distinct cell-states. A) A schematic depicting how the aggresome forms. When polyubiquitinated proteins accumulate, they are trafficked along microtubules to the centrosome. B-C) Proliferating, senescent, quiescent, and immortalized fibroblasts were treated with 0.1% DMSO or 10 μM MG132 for 8 hours prior to fixation and immunostaining for K48 polyUb (green), vimentin (red) and nuclei (Hoechst; blue). Samples were imaged, and subsequently analyzed for the proportion of cells forming aggresomes discernable by an enrichment of K48 polyUb only (red bar), a vimentin cage only (green), both (blue) or neither (white) (N=3; Two-way ANOVA with post-hoc Tukey’s test; mean ± SD). D) Examples of vimentin cages surrounding the aggresome in quiescent and immortalized fibroblasts treated with 10 μM MG132 for 8 hours prior to being fixed and immunostained as in B-C. E-F) Quiescent and immortalized fibroblasts were treated with 10 μM MG132 for 8 hours with either 0.1% DMSO or 2 mM nocodazole prior to being fixed and immunostained for K48 polyUb (green) and stained for nuclei (Hoechst; blue). Samples were quantified as in 1C. (N=3; Two-way ANOVA with post-hoc Tukey’s test; mean ± SD). G) Proliferating, senescent, quiescent and immortalized fibroblasts were treated with 10 μM MG132 for 8 hours prior to being fixed and immunostained for K48 polyUb (green), γ-tubulin (red) and stained for nuclei (Hoechst; blue). Arrows denote aggresomes (B, D, E) or centrosomes (G). Scale bars, 10 μm.

Although substantial advances have been made in understanding how aggresomes form and function, most of what is known about the aggresome comes from studies of immortalized cell lines *in vitro*. It remains unclear whether immortalization influences aggresome formation, and whether aggresomes are used by primary cell types across the body and in what cell-states. To address these gaps, we investigated aggresome formation in primary adult mouse dermal fibroblasts isolated from the tail of mice *in vitro* in four cell states: proliferating, quiescent, senescent, and immortalized. Using this model, we learned that aggresome formation in fibroblasts is dependent on immortalization and is a cell-state-specific phenotype. Despite vimentin cage formation being cited as a common feature of the aggresome and the high expression of vimentin in fibroblasts, adult fibroblasts formed the aggresome independent of vimentin cage formation (Johnston, Ward et al. 1998, Johnston, Illing et al. 2002, Morrow, Porter et al. 2020). Further, we find that the aggresome does not simply correlate with a specific type of proteostasis network, but rather that aggresomes can be employed by cells irrelevant of their preferential degradation network. Using fibroblast cell-states as a model, we performed RNA sequencing to identify features associated with the presence or absence of aggresome formation. We then validated this resource by demonstrating a role for stress-activated MAPK signaling in aggresome biogenesis, by both reproducing previous results obtained in other cell types and identifying TAK1 as a novel kinase involved in aggresome formation. Together, we present a resource which will further aid understanding of how the aggresome forms and functions, and more generally, how cell-state influences the response to proteasome inhibition.

## Results

### Aggresome formation in primary fibroblasts is cell-state specific

To determine whether primary mouse dermal fibroblasts would form the aggresome, we isolated fibroblasts from the tail of three 1 month-old C57BL/6J male mice, and then in a proliferating condition, challenged the fibroblasts with the proteasome inhibitor MG132 to induce aggresome formation (Johnston, Illing et al. 2002, Khan and Gasser 2016, Morrow, Porter et al. 2020). Cells were immunostained for two markers of the aggresome: 1) K48-linked ubiquitin (K48 polyUb), which marks proteins destined for degradation that are trafficked to the aggresome, and 2) vimentin, which forms a cage around the aggresome (Fig. 1B-C) (Johnston, Ward et al. 1998, Johnston, Illing et al. 2002, Hao, Nanduri et al. 2013). Surprisingly, although we observed a global increase in K48 polyUb, we were not able to detect an obvious enrichment of K48 polyUb at the centrosome of proliferating fibroblasts indicative of the aggresome, and similarly were not able to observe any collapse of vimentin to form a perinuclear cage, despite high levels of vimentin protein (Fig. 1B). This finding was consistent regardless of the dose and treatment length with MG132, even as cells began to undergo apoptosis (data not shown). These findings suggest that proliferating primary young adult fibroblasts do not retain the capacity to form the aggresome in response to proteasome inhibition.

Previous reports have demonstrated the formation of aggresomes in immortalized mouse embryonic fibroblast (MEF) cell lines (Lee, Nagano et al. 2010, Fusco, Micale et al. 2012, Matsumoto, Inobe et al. 2018). Therefore, we hypothesized that the aggresome may be used by fibroblasts specific to their cell-state. To test this hypothesis, we used proliferating primary young adult mouse fibroblasts to generate three additional fibroblast cell-states: 1) quiescent (contact-inhibited), 2) senescent (replicative senescence) and 3) immortalized (ectopic expression of the SV40T antigen) (Neufeld, Ripley et al. 1987, Hutter, Unterluggauer et al. 2002, Legesse-Miller, Raitman et al. 2012) (Fig. S1A-H). All cell-states were generated from the same originating pool of adult fibroblasts. Interestingly, in response to MG132, senescent fibroblasts behaved similarly to proliferating fibroblasts, exhibiting increased global K48 polyUb without forming aggresomes. However, similar to a previous report in immortalized MEFs (Lee, Nagano et al. 2010, Fusco, Micale et al. 2012, Matsumoto, Inobe et al. 2018), immortalized young adult primary mouse fibroblasts were able to form aggresomes, enriching K48 polyUb at the centrosome. Further, quiescent fibroblasts also were able to form aggresomes (Fig. 1B-C). These data suggest that change in cell-state alone is sufficient to drive an aggresome program in fibroblasts in response to impaired proteostasis. Interestingly, in both the immortalized and quiescent fibroblast populations, whereas they did form a K48 polyUb-rich aggresome, only a small proportion of cells (1.9 ± 0.49% of quiescent fibroblasts and 1.04 ± 0.82% of immortalized fibroblasts) formed collapsed vimentin cages around the aggresome (Fig. 1B-D). Taken together, these results demonstrate that the aggresome can be used by cells cell-state specifically and further suggest a decoupling of vimentin cage formation and aggresome formation, consistent with a previous study (Morrow, Porter et al. 2020).

Initially described in 1998 by Ron Kopito’s laboratory, the aggresome assembles as dynein motor proteins traffic proteins destined for degradation along microtubules to the centrosome (Johnston, Ward et al. 1998, Johnston, Illing et al. 2002, Kawaguchi, Kovacs et al. 2003, Olzmann, Li et al. 2008). Thus, aggresomes should colocalize with centrosome markers and be sensitive to microtubule depolymerization (Johnston, Ward et al. 1998). To confirm the K48 polyUb-rich structures in quiescent and immortalized fibroblasts were aggresomes, we first immunostained for γ-tubulin, a marker of the centrosome, together with K48 polyUb, confirming the perinuclear enrichments of K48 polyUb were localized to the centrosome in both quiescent and immortalized fibroblasts (Fig. 1G). We next challenged quiescent and immortalized fibroblasts with MG132 to induce aggresome formation in the presence of the microtubule poison nocodazole to block aggresome formation (Fig. 1E-F). Nocodazole efficiently blocked perinuclear enrichment of K48 polyUb in both cell-states, further confirming primary adult mouse dermal fibroblasts make aggresomes dependent upon their cell-state.

### Aggresomes are used by cells with diverse preferential proteostasis networks

Among a host of other proposed functions, the aggresome is predominantly thought to be a staging ground for protein degradation by autophagy (Iwata, Riley et al. 2005, Olzmann, Li et al. 2008). However, recent evidence suggests that additional components of the cell’s proteostasis network, such as proteasomes, also may play a role at the aggresome (Hao, Nanduri et al. 2013, Morrow, Porter et al. 2020). To gain more insight into how the aggresome is used in combination with other components of the cell’s proteostasis network and determine if the proteostasis network preferred by the cell could predict aggresome formation, we profiled the proteostasis network in each fibroblast cell-state by measuring relative levels of protein synthesis, chaperone protein expression, proteasome activity, and autophagy. First, we treated proliferating, senescent, quiescent, and immortalized fibroblasts with puromycin and measured relative levels of puromycin incorporation into nascent proteins by immunoblotting (Fig. 2A-B, S2A-B). Interestingly, we observed that whereas proliferating, senescent, and quiescent fibroblasts produced protein at relatively similar rates relative to protein content, immortalized fibroblasts exhibited significantly reduced protein synthesis compared to the other three cell-states.

**Figure 2.**
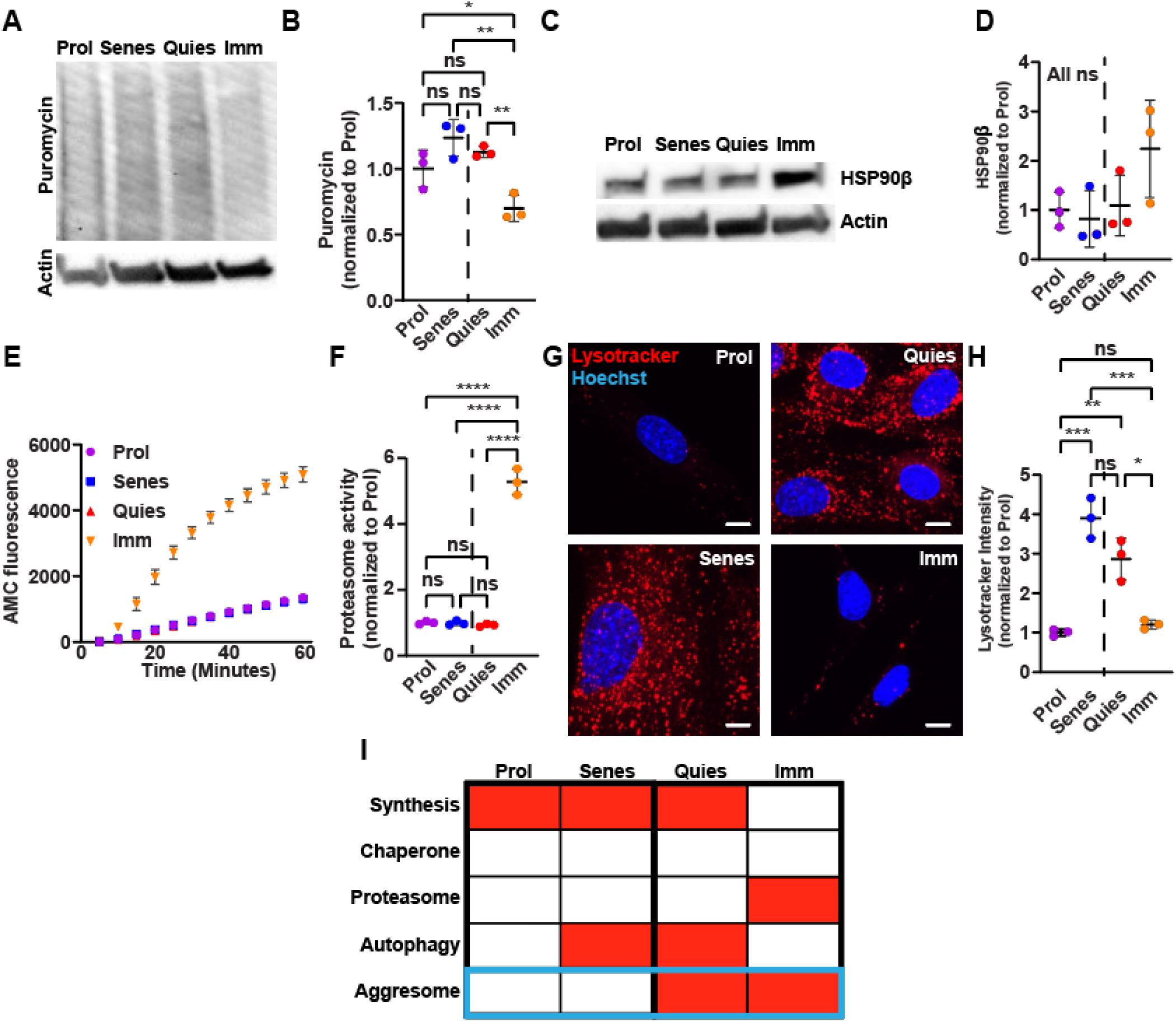
Aggresomes are used within diverse proteostasis networks. A-B) Proliferating, senescent, quiescent and immortalized fibroblasts were pulsed with 10 μg/mL puromycin for 10 minutes prior to soluble protein extraction and analysis by western blot. Relative levels of puromycin incorporation and actin expression were visualized by western blot. Samples were analyzed for puromycin levels relative to actin (N=3; Two-way ANOVA with post-hoc Tukey’s test; mean ± SD). C-D) Proliferating, senescent, quiescent and immortalized fibroblasts were analyzed for soluble protein expression of HSP90β and actin by western blot (N=3; Two-way ANOVA with post-hoc Tukey’s test; mean ± SD). E-F) Proliferating, senescent, quiescent and immortalized fibroblast protein lysates were prepared and analyzed for relative levels of trypsin-like proteasome activity by measuring AMC fluorescence as a function of time (N=3; Two-way ANOVA with post-hoc Tukey’s test; mean ± SD). F displays normalized AMC accumulation at 60 minutes for each sample. G-H) Proliferating, senescent, quiescent and immortalized fibroblasts were stained and analyzed for lysosome content (Lysotracker; red) and nuclei (Hoechst; blue) (N=3; Two-way ANOVA with post-hoc Tukey’s test; mean ± SD). The dotted line in each panel separates cell-states that either do or do not form the aggresome. I) Schematic summarizing the proteostasis network experiments in each fibroblast cell-state. Red indicates higher levels of each node of the proteostasis network respectively. Scale bars, 10 μm. *p<0.05, **p<0.01, ***p<0.001, ****p<0.0001.

Previous studies have shown that chaperone protein activity, as measured by expression of the heat shock transcription factor HSF1, was not different between proliferating human fibroblasts and senescent human fibroblasts (Sabath, Levy-Adam et al. 2020). Therefore, we hypothesized that fibroblast cell-states would not significantly differentially regulate chaperone proteins. To test this hypothesis, we probed the chaperone protein network by immunoblotting for the chaperone protein HSP90β, representing one of the five major classes of heat shock proteins (Hartl, Bracher et al. 2011, Yerbury, Ooi et al. 2016). In line with previous reports, proliferating, senescent, and quiescent fibroblasts did not display differences in protein levels of HSP90β (Fig. 2C-D). We observed trends towards increased expression of HSP90β in immortalized fibroblasts compared against the other cell-states, however these trends were not significant.

To degrade proteins, cells have two primary protein degradation systems: the ubiquitin-proteasome system and autophagy. To determine whether fibroblast cell-state drove differences in protein degradation by the ubiquitin-proteasome system, we isolated protein lysates from each fibroblast cell state, and measured proteasome activity in cell lysates generated from each cell-state by adding the substrate Boc-Leu-Arg-Arg-AMC, a molecule that becomes fluorescent when cleaved by the trypsin-like activity of the proteasome. Consistent with previous reports, proteasome activity in proliferating, senescent, and quiescent fibroblasts was similar between these cell-states, whereas immortalized fibroblasts had significantly elevated levels of proteasome activity (Legesse-Miller, Raitman et al. 2012) (Fig. 2E-F, S2C-D).

Lastly, we measured relative levels of autophagy by labeling fibroblasts with the dye Lysotracker Red DND-99. Previous data suggest that senescent and quiescent fibroblasts have relatively higher levels of autophagy compared to proliferating fibroblasts (Legesse-Miller, Raitman et al. 2012, Sun, Zheng et al. 2018, Rajendran, Alzahrani et al. 2019). Our data confirms these previous reports, as we observed that quiescent and senescent fibroblasts harbored higher levels of lysosomes compared to proliferating and immortalized fibroblasts (Fig. 2G-H, S2E-F, (Jia, Xue et al. 2018)). Taken together, fibroblast cell-states use a diverse set of proteostasis networks, and no one feature of the proteostasis network correlates with aggresome formation in fibroblasts (Fig. 2I). These data support the idea that the aggresome is used by cells in a diverse set of contexts to maintain cell fitness and that the aggresome’s mode of action is more complex than degrading proteins through autophagy.

### Transcriptomic profiling of the fibroblast response to MG132 reveals features associated with aggresome formation

We next used our fibroblast cell-state specific aggresome formation model to identify the potential molecular mechanisms promoting aggresome formation. To this end, we generated proliferating, senescent, quiescent, and immortalized fibroblasts from the same initial fibroblast population, treated with either DMSO or MG132, extracted RNA, and submitted samples for RNA sequencing to analyze transcriptomic features associated with aggresome formation (Table S1; GSE176107). To control for variance in global transcription rate across fibroblast cell states, we additionally performed an RNA spike-in prior to RNA extraction, normalized to total cell number, and then normalized gene expression to the spike-in controls prior to downstream analysis (Fig. S3A-B) (Chen, Hu et al. 2015). Multidimensional scaling (MDS) of the transcriptomic data revealed clustering across both cell-states and treatments, suggesting MG132 similarly shifted the cell’s transcriptional program, regardless of cell state (Fig. 3A). Further each cell-state clustered individually despite being genetically identical, suggesting each cell-state harbored a distinct transcriptional profile, regardless of exposure to MG132. We next asked if the proteostasis networks of cells responding to MG132 would be similar or different in cells with or without aggresomes by performing MDS on only the differential expression of genes in the proteostasis network (chaperone, proteasome, autophagy) in response to MG132 (Fig. 3B). We observed that each cell-state harbored a distinct proteostasis network transcriptional response to proteasome inhibition (Fig. 3B). Further, we asked how each node of the proteostasis network reacted in response to proteasome inhibition in each cell-state. We generated heat maps and measured the average fold change of chaperone (GO: Chaperone-mediated protein folding), proteasome (GO: Proteasome Complex), and autophagy (GO: Autophagy) genes (Fig. 3C-D). We observed stronger upregulation of chaperone and proteasome genes in response to proteasome inhibition, though autophagic machineries also became upregulated in all four cell-states (Fig 3D). Surprisingly, no one feature clearly correlated with aggresome presence, further suggesting that the differences driving aggresome formation in each cell-state are lesser than the differences driving each cell-state’s identity.

**Figure 3.**
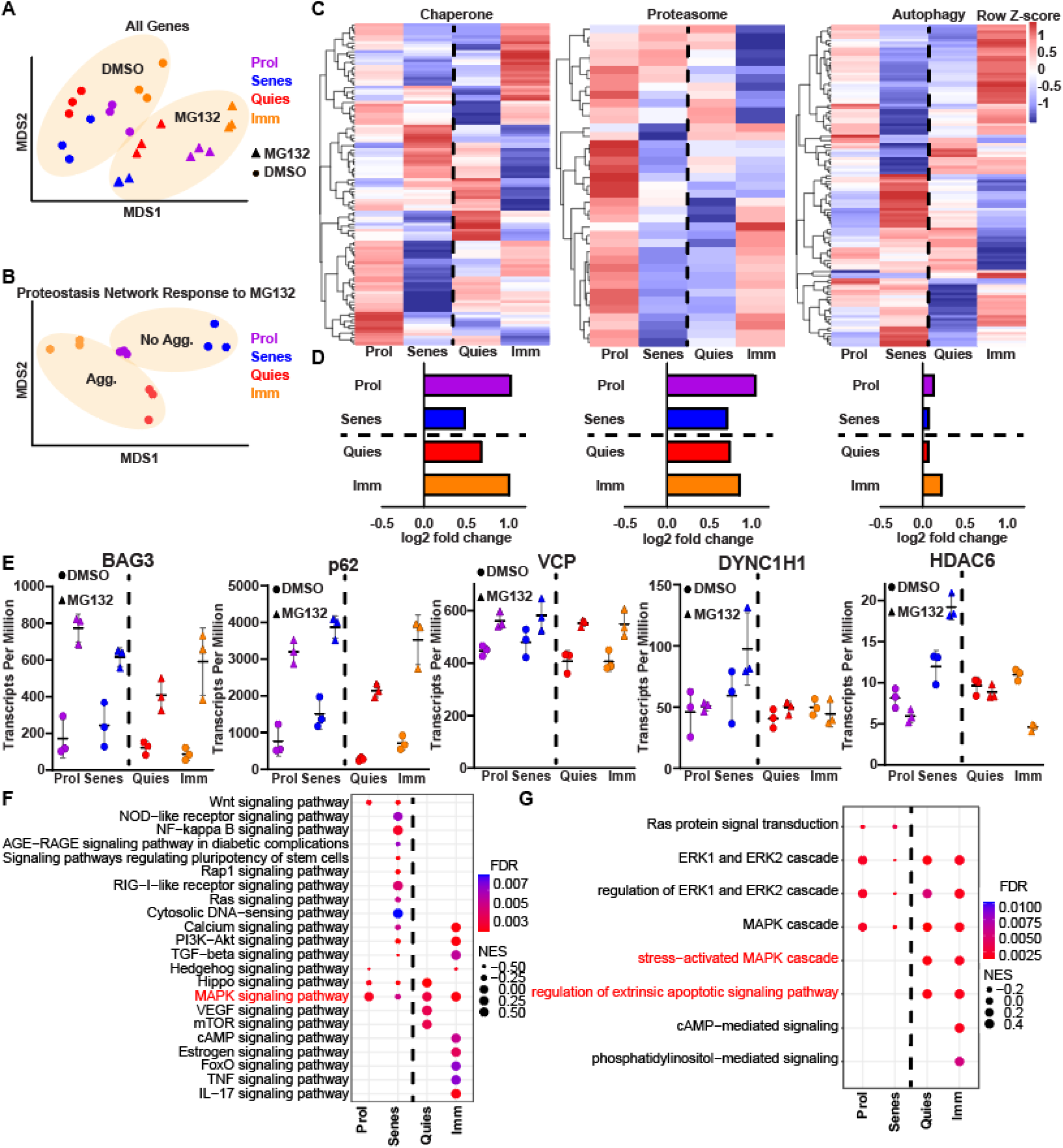
Transcriptional profiling of the fibroblast cell-state specific response to proteasome inhibition reveals stress-activated MAPK signaling associated with aggresome formation. A) MDS analysis of the global transcriptomes of proliferating (purple), senescent (blue), quiescent (red) and immortalized (orange) fibroblasts treated with either 0.1% DMSO (circles) or 10 μM MG132 (squares) for 8 hours. Ovals denote clustering of DMSO or MG132 conditions. B) MDS analysis of the proteostasis network’s (chaperone, proteasome and lysosome genes) transcriptional response to proteasome inhibition in proliferating, senescent, quiescent and immortalized fibroblasts. Ovals denote clustering of aggresome forming or lacking cell-states. C) Heat maps depicting differential expression of chaperone, proteasome, or autophagy genes in response to MG132 between proliferating, senescent, quiescent, or immortalized fibroblasts. D) Average DEseq2 log2-fold change values for all chaperone, proteasome, or autophagy genes as proliferating, senescent, quiescent and immortalized fibroblasts respond to MG132 treatment. E) Transcript per million counts of representative genes implicated in aggresome formation in proliferating, senescent, quiescent and immortalized fibroblasts treated with DMSO (circles) or MG132 (squares). F-G) GO and KEGG analysis of the signaling pathways significantly differentially regulated in response to MG132 in proliferating, senescent, quiescent, and immortalized cells. FDR refers to false discovery rate. The dotted line in each panel separates cell-states that either do or do not form the aggresome.

To gain insight into why some fibroblast cell-states form the aggresome and some do not, we reasoned that one explanation could be that genes essential for aggresome formation were downregulated or absent. Therefore, we examined the expression of known genes essential for aggresome formation or genes known to function at the aggresome, such as *SQSTM1* and *HDAC6* (Fig. 3E, Table S2). We observed expression of all known aggresome-related genes we examined in all cell-states. While it is likely that there are additional factors essential for aggresome formation that have yet to be revealed, this finding suggests that aggresome presence or absence across cell-states is not due to the lack of components to form the aggresome, but rather in how each cell-state responds to disrupted proteostasis.

Therefore, we next hypothesized that cells which form or do not form the aggresome differ in how they use their systems to respond to proteasome inhibition. To identify which features and signaling pathways would be associated with the presence or absence of the aggresome, we performed Gene Ontology (GO) and Kyoto Encyclopedia of Genes and Genomes (KEGG) analyses on each fibroblast cell-state’s transcriptional response to proteasome inhibition individually, and probed for signaling pathways unique to aggresome-forming cells (Table S3) (Fig. 3F-G). We observed that the aggresome-forming quiescent and immortalized fibroblasts uniquely significantly differentially regulated the stress-activated node of MAPK signaling and extrinsic apoptotic signaling (Fig. 3F). Therefore, we hypothesized that these signaling pathways may play a role in aggresome formation.

### TAK1 and p38 MAPK inhibition suppresses aggresome formation in immortalized and quiescent fibroblasts

To validate the predictive power of our data set in identifying signaling pathways involved in aggresome formation, we further evaluated the hypothesis that stress-activated MAPK signaling plays a role in aggresome formation in adult mouse dermal fibroblasts. Supporting our pathway analyses and hypothesis, we calculated the average log fold change of genes in the MAPK pathway generally, and more specifically in the stress-activated MAPK gene node, and found that in both cases, aggresome-forming fibroblasts (quiescent and immortalized) exhibited the strongest induction of MAPK/stress-activated MAPK signaling (Fig. 4A-B; Table S4). This observation also confirms studies that revealed an essential role for several p38 MAPKs (stress-induced MAPKs) in aggresome formation in primary hepatocytes and HEK293 cells (Nan, Dedes et al. 2006, Zhang, Gao et al. 2018, Qin, Jiang et al. 2019). Therefore, we hypothesized that p38 MAPK genes would similarly drive aggresome formation in quiescent and immortalized primary young adult mouse dermal fibroblasts.

**Figure 4.**
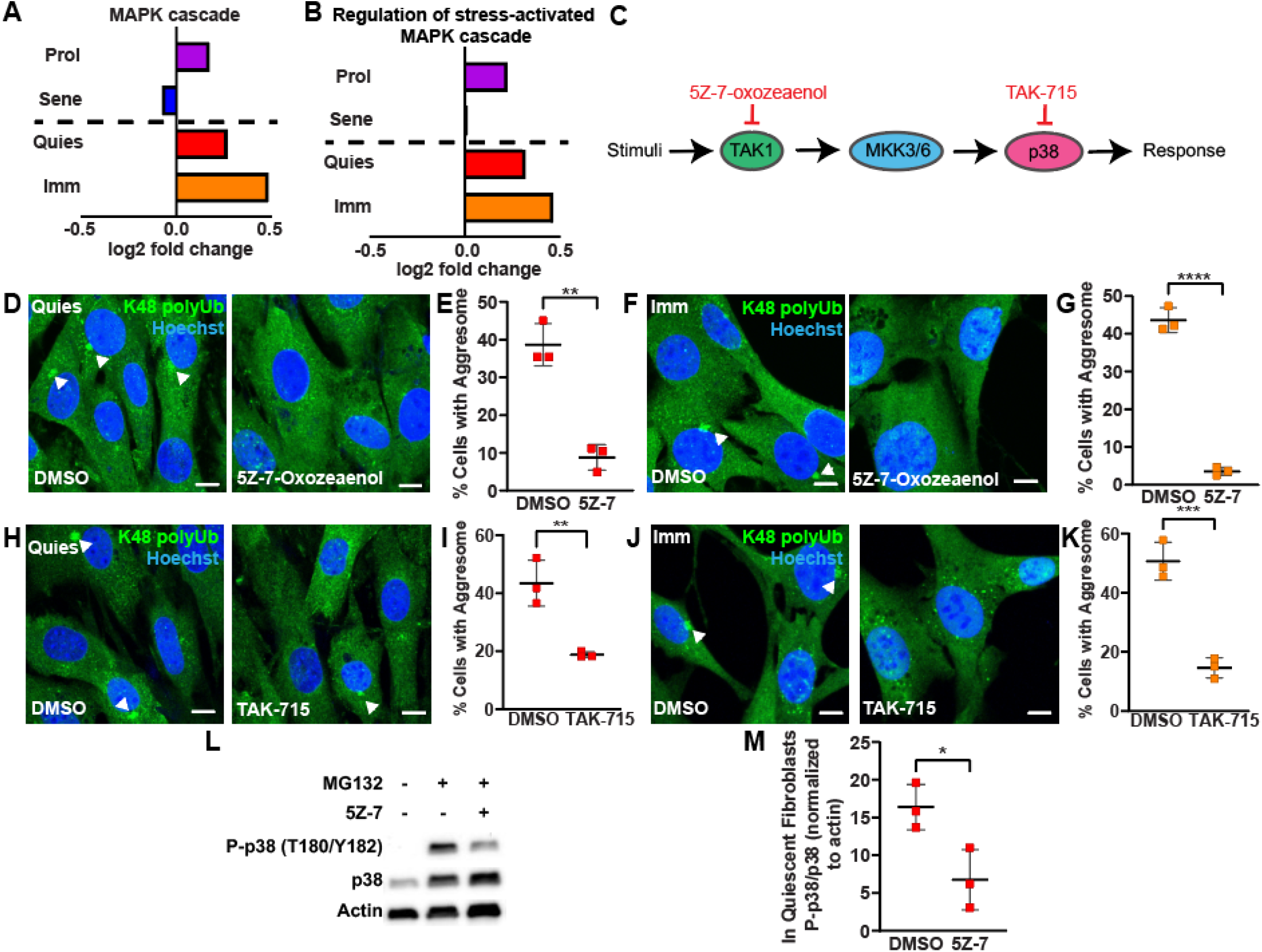
p38α MAPK and p38β MAPK inhibition with TAK-715 suppresses aggresome formation in fibroblasts. A-B) Average DESeq2 log2(fold change) values for all MAPK cascade or Regulation of stress-activated MAPK cascade genes as proliferating (purple), senescent (blue), quiescent (red) and immortalized (orange) fibroblasts respond to MG132 treatment. C) A schematic summarizing the interaction of TAK1, MKK3/6 and p38 MAPKs in stress activated MAPK signaling. Inhibitors are shown in red. D-G) Quiescent (D-E) and immortalized (F-G) fibroblasts were treated with 5 μg/mL 5Z-7-Oxozeaenol or 0.05% DMSO for 24 hours, and then additionally treated with 10 μM MG132 for 7 hours still in the presence of 5Z-7-Oxozeaenol or DMSO prior to being immunostained for K48 polyUb (green) and Hoechst (nucleus; blue). Samples were analyzed for the proportion of cells forming the aggresome (N=3; Student’s t-test; mean ± SD). H-K) Quiescent (H-I) and immortalized (J-K) fibroblasts were treated with 50 μM TAK-715 or 1% DMSO for 24 hours, then additionally treated with 10 μM MG132 for 7 hours still in the presence of TAK-715 or DMSO prior to being immunostained for K48 polyUb (green) and Hoechst (nucleus; blue). Samples were analyzed for the proportion of cells forming the aggresome (N=3; Student’s t-test; mean ± SD). L-M) Quiescent fibroblasts were treated with 10 μM MG132 and/or 5 μg/mL 5Z-7-Oxozeaenol for 8 hours as indicated in panel L and then analyzed for expression p38 MAPKs, actin and P-p38 MAPKs (T180/Y182) by western blot. Scale bars, 10 μm. *p<0.05, **p<0.01, ***p<0.001, ****p<0.0001.

To test this hypothesis and to confirm previous studies which demonstrated a role for p38 MAPKs in aggresome formation, we measured aggresome formation in quiescent and immortalized fibroblasts in the presence of a control dose of DMSO or inhibitors targeting different components of the stress-activated MAPK signaling (Fig. 4C). First, we took advantage of the TAK1 inhibitor 5Z-7-oxozeaenol (5Z-7), to determine whether inhibiting TAK1, a MAPKKK involved in stress-activated MAPK signaling, would modulate aggresome formation. We found that 5Z-7 treatment potently suppressed aggresome formation, suggesting that TAK1 is functionally important for aggresome formation. (Fig. 4D-G). This result validates the power of our transcriptomic data to provide insight into mechanisms governing aggresome formation and reveals TAK1 as a novel regulator of aggresome biogenesis.

To further validate a role for stress-activated MAPK signaling in aggresome formation, we induced aggresome formation in quiescent and immortalized fibroblasts in the presence of a control dose of DMSO or a different stress-activated MAPK inhibitor, TAK-715, an inhibitor of p38 MAPK α/β, which are thought to function downstream of TAK1. In agreement with our hypothesis and previous reports, we observed that TAK-715 treatment was sufficient to decrease aggresome formation in both quiescent and immortalized fibroblasts (Fig. 4H-K) (Qin, Jiang et al. 2019). This finding further validates a role for stress-activated MAPK signaling in aggresome formation. As TAK1 is thought to act upstream of p38 MAPKs, we next hypothesized that TAK1 could be driving aggresome formation through p38 MAPKs. To test this hypothesis, we measured phosphorylation of p38 MAPKs at T180/Y182, the site of TAK1-mediated phosphorylation through MKK3/6, in quiescent fibroblasts challenged with MG132 and either a control dose of DMSO or the TAK1 inhibitor 5Z-7. We observed that 5Z-7 reduced phosphorylation of p38 MAPK at T180/Y182, suggesting that TAK1 drives aggresome formation, at least in part, through driving phosphorylation of p38 MAPKs (Fig. 4L-M). These data support a role for stress-activated MAPK signaling in aggresome formation in adult fibroblasts and reveal the utility of our model system whereby genetically identical adult primary cells in the same media conditions and without overexpression of proteotoxic factors, differing only in cell-state, can be dissected for drivers of aggresome formation.

Here we demonstrate that primary young adult mouse dermal fibroblasts use the aggresome in the quiescent and immortalized cell-state, but not in the proliferating and senescent cell-state. We find that aggresome formation is not a reflection of a cell’s preference for protein degradation by autophagy, but rather, that each fibroblast cell-state employs a unique proteostasis network to maintain proteostasis. Transcriptomic analysis of the fibroblast response to proteasome inhibition revealed features associated with cells that form aggresomes. Among these features, we found stronger activation of stress-activated MAPK signaling is associated with aggresome formation, supporting studies showing p38 MAPKs drive aggresome formation in other cell types (Nan, Dedes et al. 2006, Zhang, Gao et al. 2018, Qin, Jiang et al. 2019, Magupalli, Negro et al. 2020). Confirming these data, we found that inhibition of both TAK1 and p38α/β MAPKs was sufficient to suppress aggresome formation in fibroblasts. Taken together, we demonstrate that the aggresome formation is not a universal component of a cell’s response to disrupted proteostasis, but rather that aggresomes can be used cell-state specifically, and provide resources for a better understanding of aggresome formation and function.

## Discussion

Here we demonstrate that preference to degrade protein through the aggresome is cell-state specific (Fig. 1). However, it is unclear whether the absence of aggresome formation is a reflection of proliferating and senescent fibroblasts making a decision to not form the aggresome through suppression of a pathway, compared to an inability to form the aggresome due to missing expression of unknown aggresome formation machineries. Regardless, if the aggresome is cytoprotective as evidence suggests (Tanaka, Kim et al. 2004), it is not clear why some cells would not evolve to use this mechanism for maintaining proteostasis, while other cells have conserved this mechanism. One possibility could be that redundancy in the cell’s proteostasis network allowed for cell types to evolve divergently in how they maintain proteostasis. It also could be that we still don’t fully understand the function of the aggresome and that there may be more nuanced contexts which dictate when the aggresome would be the most useful for a cell’s fitness.

These data also provide an important cautionary note regarding the use of immortalized cell lines to study biology that is reflective of cell behavior *in vivo*. The majority of aggresome literature to date has investigated the aggresome in immortalized cells *in vitro*. However, our data demonstrate that the *simple* act of immortalizing cells with SV40T is sufficient to drive cells that would not otherwise do so, to use the aggresome (Fig. 1). Additionally, cell immortalization drove widespread remodeling of the proteostasis network (Fig. 1). While initial studies of the aggresome in immortalized cell-lines *in vitro* have provided great utility towards an understanding of aggresome function and mechanisms of aggresome formation and should not be discounted, our data highlights the importance of translating immortalized cell line findings to primary cell lines *in vitro* and to cells in tissues *in vivo*.

In our report, we identify TAK1 as a novel regulator of aggresome formation acting at least in part through driving phosphorylation and activation of p38 α/β MAPKs (Fig. 4). As TAK1 can be activated by TGF-β, identification of TAK1 as a mediator of aggresome formation is timely, as recent work has described a role for aggresome-like structures as a program implicated in the cell’s inflammasome (Magupalli, Negro et al. 2020). When specific types of inflammasomes form, an HDAC6-dependent perinuclear puncta of proteins implicated in the cell’s response to infection assembles. Thus, our data provide a mechanism linking cytokine signaling and formation of the aggresome, which could corroborate a role for the aggresome in the cell’s immune system. Further, TAK1 has many targets outside of p38 MAPKs which could additionally play a role in aggresome formation such as c-Jun N-terminal kinases (JNKs) and Nuclear Factor kappa-light-chain-enhancer of activated B cells (NF-kB) (Totzke, Scarneo et al. 2020). Lastly, although TAK-715 and 5Z-7-oxozeaenol are recognized as inhibitors of p38α and p38β MAPKs or TAK1 respectively (Ninomiya-Tsuji, Kajino et al. 2003, Miwatashi, Arikawa et al. 2005, Verkaar, van der Doelen et al. 2011, Takano, Uchida et al. 2019), both of these drugs are known to have off-target effects. Therefore, it is possible that the effects we observe from these drugs on aggresome formation is through their effect on other proteins.

Lastly, while our model allows us to do many comparative analyses to understand aggresome formation and function, the heterogeneous response within each cell-state (e.g. quiescent cells treated with MG132 will maximally only be 30-50% aggresome+) leads to difficulties in interpreting bulk analyses such as western blots, versus single cell analyses such as immunofluorescence. Further, primary young adult mouse dermal fibroblasts are very resistant to genetic manipulation, rendering further detailed knockout experiments unfeasible. Future studies in amenable primary cell types will be needed to unravel the intricate relationships within these pathways.

In summary, we identify primary young adult mouse dermal fibroblast cell-states as a model to understand aggresome formation and function, and further use this system to reveal the aggresome as a non-universal, but tunable mechanism cells can activate in a cell-state specific manner to maintain proteostasis through multiple mechanisms, specifically through stress-activated MAPK signaling.

## Materials and Methods

### Fibroblast Isolation, Cell-State Generation, and Culture

Following a previously established protocol, fibroblasts were taken from tail snips of male 4 week-old C57Bl/6J mice (Khan and Gasser 2016). To isolate the fibroblasts for *in vitro* culturing, tail snips were incubated in 70% ethanol for 5 minutes at room temperature. Tail snips were then cut into smaller pieces with scissors and put into an enzyme solution (20 μM Protease (Sigma P8811), 35 μM Collagenase D (Sigma 11088866001), 1.5 mM Tris Buffer and 62.5 μM EDTA) to dissociate the tissue and release cells at 37°C for 90 minutes. Samples were then mashed for 2 minutes with a plunger from a syringe and then filtered through a 70 μm cell strainer. The sample was then washed by adding 10 mL of fibroblast culture media (FCM; RPMI with Glutamine (Invitrogen 11875-093), 10% FBS (Invitrogen 16000044), 55 μM β-Mercaptoethanol (Invitrogen 21985-023), 100 μM asparagine and 100 μg/ml Penicillin-Streptomycin (Invitrogen 15140122), 25 μg/mL Amphoterecin B (Sigma-Aldrich A9528)) and cells were pelleted at ~580 x g for 7 minutes. Cells were washed a second time with 10 mL of FCM and spun down similarly. Cell pellets were then resuspended and plated into one well of a 6 well plate (Fisher Scientific 1483211). Cells were left to sit for 5 days at 37°C and 5% CO_2_ after initial plating and were only fed once after three days by adding an additional volume of FCM. After 1-2 weeks fibroblasts expanded and were ready for experiments, and were fed at least once every 3 days with FCM.

Confluent fibroblasts were washed once with PBS, trypsinized with 0.25% trypsin-EDTA (Gibco 25200072) for 3-4 minutes at 37°C and 5% CO_2_, spun down at 400 x g for 4 minutes, and then resuspended and plated into FCM. To generate proliferating fibroblast cultures, fibroblasts were seeded at ~30% confluence and allowed to sit for 2-3 days prior to analysis and prior to reaching greater than 80% confluence. To generate senescent fibroblasts (β-galactosidase+, LaminB1 low), fibroblasts were plated at 10% confluence and then passaged sequentially at 10% confluence until the cultures no longer expanded. After cultures stopped expanding, all cultures were left for at least 7 days to develop a mature senescence phenotype, with regular media changes at least every 3 days. To generate quiescent fibroblasts (EdU-), fibroblasts were seeded at 70-80% confluence. After reaching 100% confluence, cultures were left for at least 3 additional days to develop a mature quiescent phenotype, and during this time still fed at least once every 3 days. To immortalize fibroblasts (SV40t+), low passage proliferating fibroblasts (P0-2) were seeded at 50% confluence and then transduced with retroviral particles harboring SV40t (as described in the “HEK Culture” and “Viral Particle Production” section). After transduction, fibroblasts were seeded at 5% confluence and then immortalized cells brought the dish to confluency. After 2 passages, the cultures were pure for immortalized cells as detected by Sv40t immunostaining. Immortalized fibroblasts were used up until passage 10.

### HEK293T Culture and Viral Particle Production

HEK293T cells (gift from Dr. Subhojit Roy) were cultured in HEK293T media (DMEM, high glucose, pyruvate, no glutamine; Thermo 10313021) supplemented with 10% FBS (Invitrogen 16000044) and Penicillin-Streptomycin (Invitrogen 15140122) at 37°C and 5% CO_2_. When 100% confluent, cells were trypsinized with TrypLE Express Enzyme (Invitrogen 12604013) for 3 minutes at room temperature and spun down at 120 x g for 4 minutes prior to being resuspended in HEK293T media and replated. To generate viral particles, HEK293T cells were transfected using lipofectamine (Invitrogen 11668019) with CMV-GP (gift from Dr. Fred Gage), pCMV-VSV-G (gift from Dr. Fred Gage) and a transfer plasmid (SV40T; gift from Dr. Peter Lewis). 48 hours after transfection, viral particles were collected in the supernatant, sterile filtered to remove HEK293T debris, and applied directly to fibroblasts.

### Microscopy

All images were taken with either a Nikon C2 confocal microscope or a Zeiss widefield epifluorescent microscope. In figures 1-4, images represent max intensity projections of z-stacks taken to capture the entire cell with 1 μm step sizes. For live-cell experiments, cells were kept in a stage-top incubator to maintain humidity, 37°C, and 5% CO_2_. Images were analyzed using Fiji.

### Immunostaining

To immunostain fibroblasts to detect vimentin (1:1,000; Sigma-Aldrich AB5733), K48 polyubiquitin (1:500; Millipore Sigma 05-1307), Lamin B1 (1:500; Abcam ab16048), SV40T (1:500; Abcam ab234426), GFP (1:500; Aves Labs GFP-1020) and fibroblasts were fixed in 4% paraformaldehyde (PFA) for 15 minutes at room temperature. Samples were then permeabilized with 0.25% triton for 15 minutes at room temperature and blocked with 20% donkey serum (Millipore Sigma S30-100ML) in an antibody buffer (150mM sodium chloride, 50mM tris base, 1% bovine serum albumin (Sigma-Aldrich A2153-50G), 100mM L-lysine, 0.04% sodium azide, pH 7.4). Primary antibodies were diluted as listed above in antibody buffer and then applied to samples overnight at 4°C. The following day, samples were washed 3 times with PBS for 10 minutes at room temperature and then stained with secondary antibodies (1:500) for 1.5 hours at room temperature. Following secondary staining, samples were washed 3 times with PBS for 10 minutes. In the second to last wash, samples were treated with Hoechst to label nuclei for 10 minutes at room temperature (1:10,000 in PBS; Thermo Fisher Scientific 62249).

To immunostain fibroblasts with γ-tubulin (1:250; Sigma-Aldrich T5326) and K48 polyubiquitin (1:500; Millipore Sigma 05-1307), cells were stained for K48 polyubiquitin as described above. Following staining for K48 polyubiquitin, cells were further stained to detect γ-tubulin by performing the following protocol. Cells were permeabilized with ice-cold methanol (pre-chilled for 1 hour at −80°C) for 10 minutes at −20°C. Samples were then blocked and stained with primary and secondary antibodies as described above.

### Western Blotting

Western blots probing for actin (1:1,000; Bio Rad VMA00048), HSP90β (1:1,000; GeneTex GTX101448), p38 MAPK (1:1,000; Cell Signaling Technology 8690L), P-p38 MAPK (1:1,000; Cell Signaling Technology 4511S) and puromycin (1:1,000; Millipore Sigma MABE343) were performed using the following protocol. To extract soluble proteins, cell pellets were resuspended in a soluble lysis buffer (10 mM Tris/HCl, 150 mM NaCL, 0.5 mM EDTA, 0.5% NP40, with protease inhibitor added) and placed on ice for 30 minutes. To extract total protein fractions, cell pellets were resuspended in total lysis buffer (10 mM Tris/HCl, 1% Triton, 150 mM NaCl, 10% Glycerol, 4% SDS, with DNAse and protease inhibitor added) and lysed for 30 minutes. Following lysis, samples were sonicated with a probe sonicator 15 times for 1 second on low power, and then clarified by centrifugation at 15,000 x g for 10 minutes. Samples were then quantified using the DC assay (Bio-Rad 5000112) and then ~5 μg of protein per sample was run through an SDS-PAGE gel. SDS-PAGE gels were transferred onto a membrane at 100V for 1.5 hours at 4°C. Membranes were then blocked in TBS-T with 5% milk for 30 minutes at room temperature. Primary antibodies were diluted in 5% milk and incubated on membranes overnight diluted as indicated above at 4°C on a tube roller. Primary antibody was washed off 3 times with TBS-T for 5 minutes/wash at room temperature on a tube roller. Secondary antibodies were applied to membranes in 5% milk in TBS-T for 1.5 hours at room temperature and then washed off 3 times with TBS-T for 5 minutes/wash on a tube roller. Following washing off secondary, protein expression was detected using a UVP Imaging system after treating blots with an ECL reaction. To quantify protein expression, blots were analyzed in ImageJ. In brief, ROIs were drawn around each band and measured, background signal was subtracted, and all proteins were normalized to housekeeping genes, or as otherwise noted in the figure legends. All experiments were repeated at least 3 times to ensure reproducibility.

### EdU Pulse

Fibroblasts were pulsed for 1 hour with 10 μM EdU (Invitrogen C10337) at 37°C to label cells in S-phase and then fixed with 4% PFA for 15 minutes at room temperature. Cells were then treated per the manufacturer’s recommendations to visualize EdU through click-chemistry (Invitrogen C10337). Cells were stained with Hoechst to label nuclei (1:10,000; Thermo Fisher Scientific 62249). Samples were then analyzed for the percentage of cells that were EdU+.

### Aggresome Assays

To detect aggresomes in various conditions, cells were treated as described in each legend respectively (MG132 – Sigma M7449; 5Z-7-oxozeaenol – Sigma O9890; TAK-715 – MedChem Express HY-10456; nocodazole – Sigma M1404). Equal volume DMSO was applied to cells as a vehicle control for compounds dissolved in DMSO. Cells were immunostained as described in the “Immunostaining” section of the methods and analyzed for the percentage of cells with an aggresome as determined by an observer blind to experimental conditions but trained to identify aggresomes. An aggresome was defined as a perinuclear enrichment of K48 polyUb localizing to the nuclear bay. All aggresome quantification experiments were repeated at least 3 times.

### B-Galactosidase Assays

Proliferating and Senescent mouse fibroblasts were washed twice with 1X PBS and fixed and stained for B-Gal activity using the Senescence B-Galactosidase Staining Kit per the manufacturer’s instructions (Cell Signaling Technology 9860S). The following day cells were washed twice with 1X PBS and stained with Hoechst (Thermo Fisher Scientific 62249). After staining, cells were imaged using a white light microscope and the percentage of B-Gal+ cells was determined by taking the number of B-Gal+ cells over the number of Hoechst+ cells in each image. This experiment was repeated 3 times.

### Proteasome Activity Assays

Protein was extracted from fibroblasts with conditions optimized to preserve proteasome activity by resuspending pellets in 50mM Tris-HCl, pH 7.5, 250mM sucrose, 5mM MgCl2, 0.5mM EDTA and 1mM dithiothreitol. Samples were then lysed by passing samples through a 27-gauge needle ten times. Samples were clarified through centrifugation for 10 minutes at 4°C at 15,000 x g. Protein concentration was quantified using the DC protein assay (Bio-Rad 5000112). To perform the assay, 10-30 μg of protein were loaded for each sample (equal weight of total protein was loaded across samples in each experiment) into a reaction buffer comprised of: 2 mM adenosine triphosphate (Fisher ICN15026605), 0.37 mM proteasome substrate (trypsin - Boc-Leu-Arg-Arg-AMC) and then diluted to 100 μl in resuspension buffer. Samples were measured every 5 minutes over the course of 1 hour in a plate reader at 37°C by exciting at 380 nm and collecting at 460 nm. Three wells were averaged for each sample and the experiment was repeated 3 times on three separate days.

### Lysotracker Assays

Live fibroblasts were treated with Lysotracker Red DND-99 (1:1,000; Thermo Fisher Scientific L7528) and Hoechst (1:10,000; Thermo Fisher Scientific 62249) for 30 minutes at 37°C and 5% CO_2_ to label lysosomes and nuclei respectively. Following dye labeling, cells were washed 3 times with media and immediately imaged. To analyze the data, z stacks were acquired, and raw integrated intensity normalized to area was calculated for each cell using ImageJ. 1 μM chloroquine was used as a positive control to validate Lysotracker. Each experiment was repeated at least 3 times.

### Protein Synthesis Assays

Fibroblasts were pulsed with 10 μg/mL puromycin (Invitrogen A11138-03) for 10 minutes at 37°C and 5% CO_2_ to label nascent proteins. As a control, samples were treated with the protein synthesis inhibitor 10 μM cycloheximide (Millipore Sigma C4859-1ML) for 10 minutes prior to puromycin treatment and during puromycin treatment. After treatment, protein was extracted and analyzed by western blot as described in the “Western Blot” section of the methods. This experiment was repeated 3 times.

### RNA extraction, library preparation and sequencing

Fibroblast cell-states were generated as described in “Fibroblast Isolation, Cell-State Generation and Culture” and then treated with either a vehicle control dose of 0.1% DMSO or 10 μM MG132 for 8 hours to induce aggresome formation. After 8 hours, cells were trypsinized and counted and then pellets of equal cell number were created. Prior to RNA extraction, ERCC Spike-In Mix RNA was added to resuspended pellets at a ratio of 0.5 μL (of 1:100 diluted from ERCC stock received from the manufacturer) to 100,000 cells to provide the capacity to normalize mRNA to cell number and probe for changes in global transcription across cell-states and treatments. Total RNA was extracted using a RNeasy Mini Kit (Qiagen 74104) according to the manufacturer’s instructions. Extracted RNA was submitted to LC Sciences for poly(A) RNA sequencing.

Poly(A) RNA sequencing libraries were generated using Illumina’s TruSeq-stranded-mRNA protocol. Poly(A) mRNAs were isolated using oligo-(dT) magnetic beads, fragmented using divalent cation buffer at elevated temperature and then used to construct DNA libraries for sequencing. Samples were quality controlled using an Agilent Technologies 2100 Bioanalyzer High Sensitivity DNA Chip and then sequenced as paired-end 140 base pair reads on an Illumina NovaSeq 6000 sequencing system.

### RNA-seq data processing

Sequencing data was trimmed using Cutadapt 1.10 (Martin 2011). Reference mouse transcripts were obtained from the Ensembl 99 build (GRCm38p6), and combined with the ERCC92 sequences, to create a common mapping reference index using kallisto 0.43.0 (Bray, Pimentel et al. 2016). Trimmed reads were mapped to the combined reference with default parameters.

Parsed pseudocount values were extracted for each sample, and data was imported into R 3.6.3 for processing. We leveraged the R package ‘RUVseq’ 1.20.0 (Risso, Ngai et al. 2014) to normalize counts as a function of the ERCC92 spike in count values using the RUVg function. RUV-normalized counts were then used as input to R package DESeq2 version 1.30.1 to perform differential expression analysis for each fibroblast cell-state’s response to proteasome inhibition. MDS analyses were performed using cmdsale() from base R and spearman rank correlation across samples in R. Heat maps were created using pheatmap() in R scaling by row. KEGG analyses was performed using gseKEGG() from the clusterprofiler V3.13 package in R with a FDR (false discovery rate) cutoff of 0.05. GO analyses were performed using gseGO() from the clusterprofiler V3.13 package in R with a FDR cutoff of 0.05. Average fold change plots were generated by taking the average differential expression output from DESeq2 for each gene in each pathway respectively. Gene lists for each node of genes came from the Gene Ontology Browser (http://www.informatics.jax.org/vocab/gene_ontology - accessed in February 2021). Transcript per million counts (TPM) were calculated by taking the raw gene counts normalized to ERCC spike-in and dividing by gene length in kilobases to get reads per kilobase (RPK) (https://www.rna-seqblog.com/rpkm-fpkm-and-tpm-clearly-explained/). Then RPK was divided by an RPK scaling factor equal to the sum of all RPK values in each sample divided by 1 million. Raw data is available online: GSE176107. Scripts for performing these analyses can be found at: https://github.com/chrismorrow5/Fibroblast-Aggresome-RNA-Sequencing-Analysis.

### Statistics

All experiments, unless otherwise noted, were repeated at least three times on three separate days with at least three technical replicates, as applicable. Typically data in the figures displays average values for technical replicates from one experiment that is representative of all experiments performed with error bars representing standard deviation. Significance tests were performed using GraphPad Prism using tests as indicated in each figure legend.

### Data Accessibility

All data and materials generated in this study are available upon request to the corresponding author (Darcie Moore). Code used is available at: https://github.com/chrismorrow5/Fibroblast-Aggresome-RNA-Sequencing-Analysis. The RNA Sequencing data is available at: GSE176107.

## Acknowledgments

We thank Peter Lewis for the SV40t construct. We thank our funding sources: NIH T32 T32GM008688 (to C.S.M.), Diana Jacobs Kalman Fellowship from AFAR (to C.S.M.), Wisconsin Graduate Fellowship (to C.S.M.), Sloan Foundation fellowship (to D.L.M.), DP2 1DP2OD025783 NIH New Innovator Award (to D.L.M.).

## Author Contributions

C.S.M. and D.L.M. conceived and designed the study. C.S.M. performed most of the experiments. Z.P.A. performed the β-galactosidase assays. B.P. performed the proteasome activity assays. E.Z. optimized the fibroblast isolation protocol and performed the initial proliferating fibroblast aggresome experiments. B.A.B. performed the RNA sequencing spike-in normalization. C.S.M. and D.L.M. wrote the manuscript with input from all authors.

---

**Figure S1.**
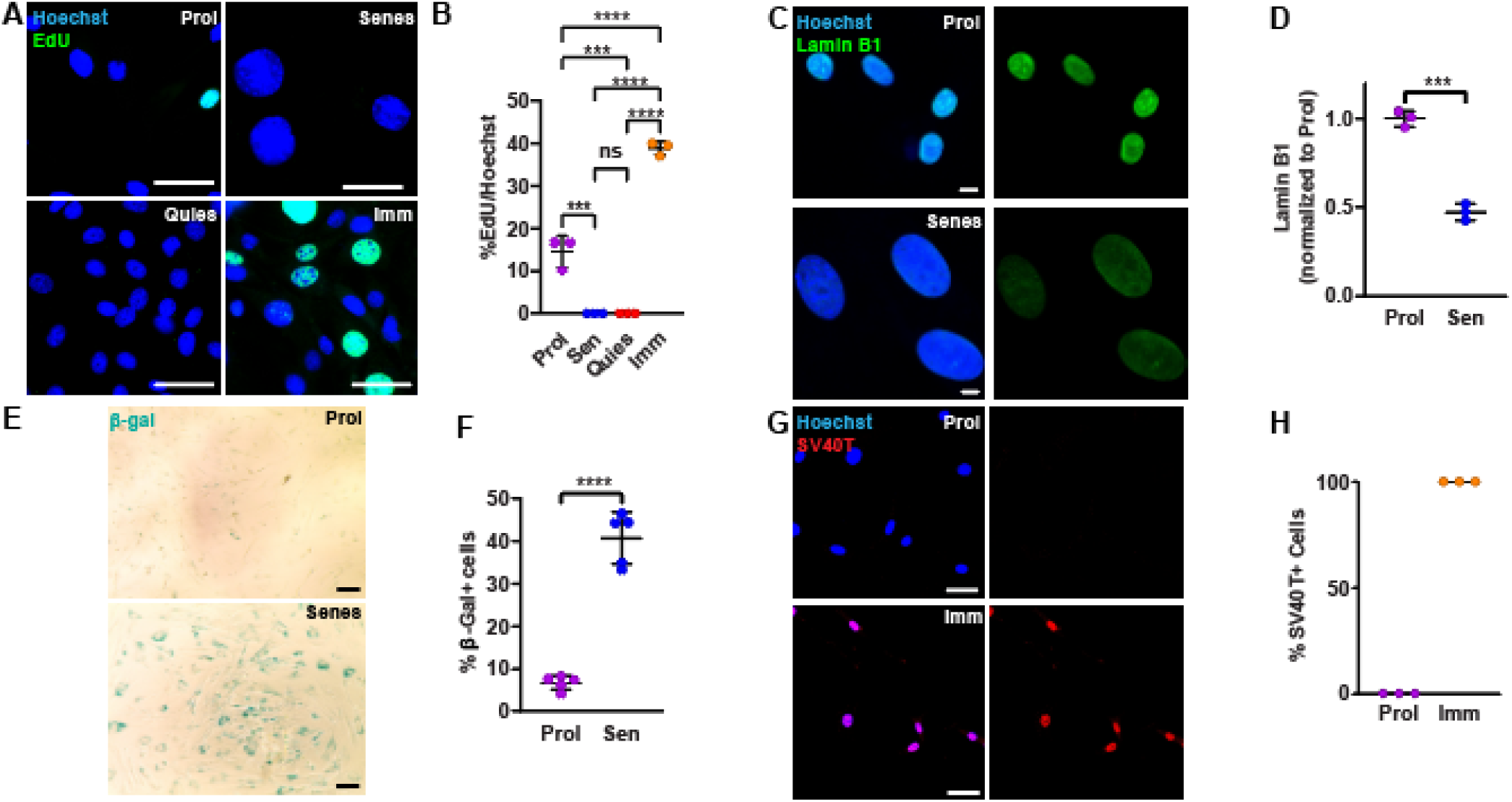
A-B) Proliferating (purple), senescent (blue), quiescent (red) and immortalized (orange) fibroblasts were pulsed with EdU for 1 hour at 37°C prior to being fixed and stained to visualize nuclei (Hoechst; blue) and EdU (green). Cells were analyzed to identify the proportion of cells that were EdU+ (N=3; Two-way ANOVA with post-hoc Tukey’s test; mean ± SD). C-D) Proliferating (purple) and senescent (blue) fibroblasts were fixed and immunostained for Lamin B1 (green) and stained for nuclei (Hoechst; blue). Samples were analyzed for relative intensity of Lamin B1 (N=3; Student’s t-test; mean ± SD). E-F) Bright field image of proliferating (purple) and senescent (blue) fibroblasts that were fixed and stained to visualize β-galactosidase activity (β-gal; green). Samples were analyzed for the proportion of cells positive for β-galactosidase activity (N=3; Student’s t-test; mean ± SD). G-H) Proliferating and immortalized fibroblasts were fixed and immunostained for SV40T (red) and stained for nuclei (Hoechst; blue). Samples were analyzed for the percentage of cells positive for SV40T (N=3; mean ± SD). Scale bars, 50 μm (A, E, G), 10 μm (C). ***p<0.001, ****p<0.0001.

**Figure S2.**
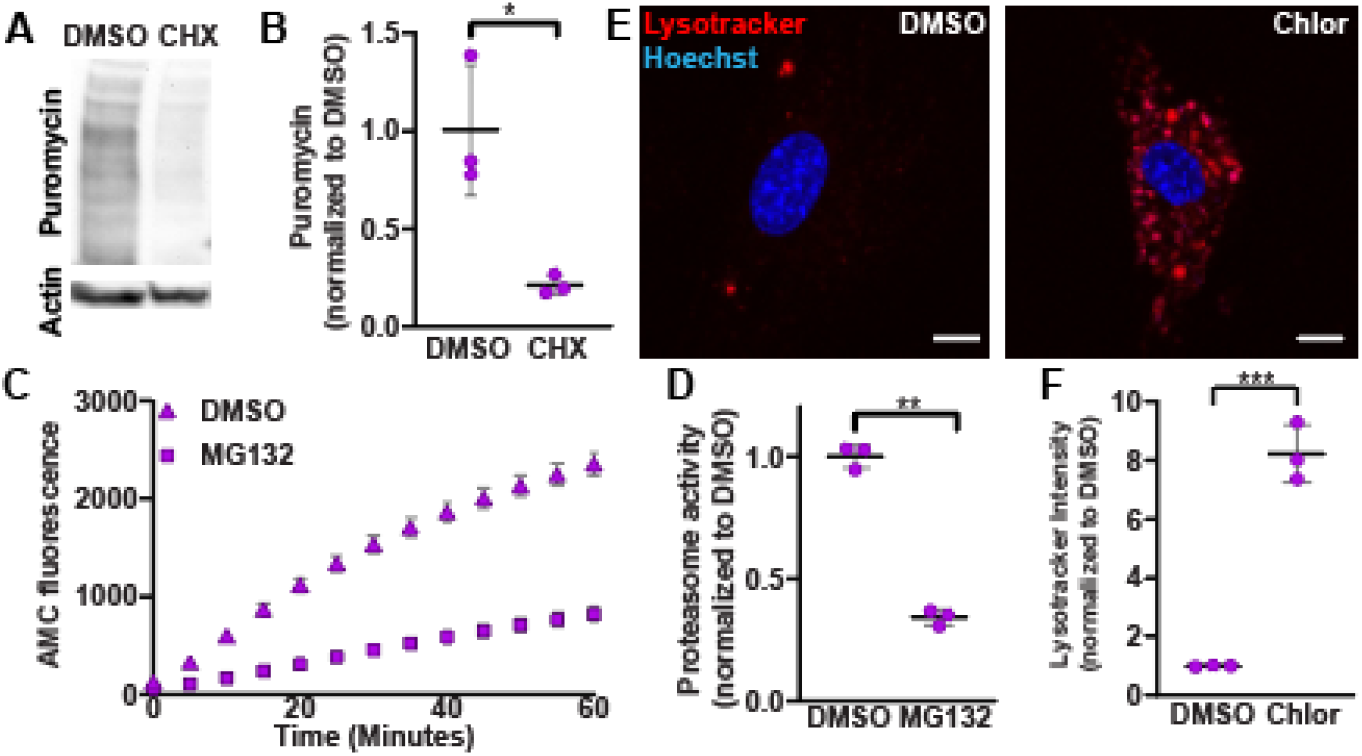
A-B) Proliferating fibroblasts were treated with 10 μM cycloheximide or 0.1% DMSO for 10 minutes prior to addition of 10 μg/mL puromycin and then protein extraction and analysis of puromycin incorporation and actin expression by western blot. Samples were analyzed for relative puromycin incorporation relative to actin (N=3; Student’s t-test; mean ± SD). C-D) Proliferating fibroblast protein lysates were prepared and treated with either a control dose of 1% DMSO or 100 μM MG132 and then analyzed for relative levels of proteasome activity by measuring AMC fluorescence as a function of time. D displays normalized AMC accumulation at 60 minutes for each sample (N=3; Student’s t-test; mean ± SD). E-F) Proliferating fibroblasts were treated with either 0.1% DMSO or 1 μM chloroquine for 24 hours and then stained and analyzed for lysosome content (Lysotracker; red) and nuclei (Hoechst; blue) (N=3; Student’s t-test; mean ± SD). Scale bars, 10 μm. *p<0.05, **p<0.01, ***p<0.001.

**Figure S3.**
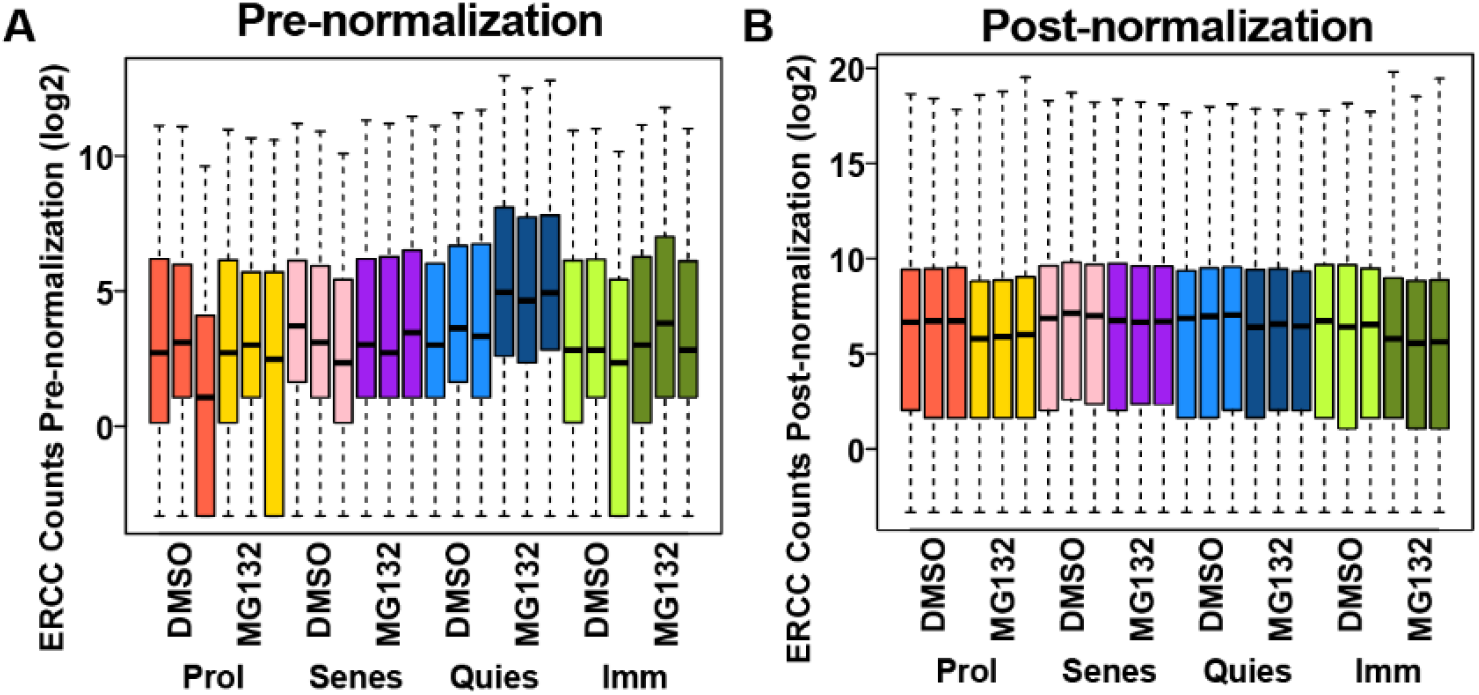
A-B) ERCC spike-in counts relative to other transcripts in proliferating (prol), senescent (senes), quiescent (quies), and immortalized (imm) fibroblasts treated with DMSO or MG132 pre-ERCC normalization (A) or post-ERCC normalization (B). All RNA sequencing analyses shown are analyses performed using the post-normalized data.

